# Estimating strength of polygenic selection with principal components analysis of spatial genetic variation

**DOI:** 10.1101/008011

**Authors:** Davide Piffer

**Keywords:** Intelligence, Natural Selection, Polygenic evolution, Principal Components Analysis

## Abstract

Principal components analysis on allele frequencies for 14 and 50 populations (from 1K Genomes and ALFRED databases) produced a factor accounting for over half of the variance, which indicates selection pressure on intelligence or genotypic IQ. Very high correlations between this factor and phenotypic IQ, educational achievement were observed (r>0.9 and r>0.8), also after partialling out GDP and the Human Development Index. Regression analysis was used to estimate a genotypic (predicted) IQ also for populations with missing data for phenotypic IQ. Socio-economic indicators (GDP and Human Development Index) failed to predict residuals, not providing evidence for the effects of environmental factors on intelligence. Another analysis revealed that the relationship between IQ and the genotypic factor was not mediated by race, implying that it exists at a finer resolution, a finding which in turn suggests selective pressures postdating sub-continental population splits.

Genotypic height and IQ were inversely correlated but this correlation was mostly mediated by race. In at least two cases (Native Americans vs East Asians and Africans vs Papuans) genetic distance inferred from evolutionarily neutral genetic markers contrasts markedly with the resemblance observed for IQ and height increasing alleles.

A principal component analysis on a random sample of 20 SNPs revealed two factors representing genetic relatedness due to migrations. However, the correlation between IQ and the intelligence PC was not mediated by them. In fact, the intelligence PC emerged as an even stronger predictor of IQ after entering the “migratory” PCs in a regression, indicating that it represents selection pressure instead of migrational effects.

Finally, some observations on the high IQ of Mongoloid people are made which lend support to the “cold winters theory” on the evolution of intelligence.

## Introduction

IQ and height are highly polygenic traits (Allen et al., 2010; Benyamin et al., 2013), which show strong heritability (Plomin et al., 2008).

These traits also show considerable variation between populations and racial groups (Lynn & Vanhanen, 2012). Intense effort has been spent on trying to identify genetic variants that account for phenotypic variation within populations. Recently, attempts have been made to explain phenotypic differences between populations using genetic variants with known effects on height (Turchin et al., 2012; Piffer, 2014) and intelligence (Piffer, 2013; Woodley et al., 2014). Encouraging results also come from a recent population genetics study of happiness, which showed that the frequency of a genetic variant (5-HTT) and Danish ancestry predict happiness among countries and between individuals even after accounting for GDP and socio-economic variables (Proto & Oswald, 2014).

The goals of this paper are: a) to find out whether the genotypic IQ factor predicts phenotypic IQ also after controlling for socio-economic indicators and genetic variation due to human migrations; b) to predict genotypic IQ scores for populations in the 1K Genomes and ALFRED data set. This is an important test of the model because specific predictions are generated that can be tested by assessing the IQ of populations whose phenotypic IQ has not been estimated yet; moreover it allows us to: c) test the environmentalist hypothesis that residuals (countries whose estimated IQs are lower or higher than predicted by their genetic factor scores) are explained by socioeconomic factors. Finally: d) to shed light on the correlation between height and intelligence at the population genetics and evolutionary level.

A relatively well-known study (Chabris et al., 2012) claimed that, although a number of candidate genes had been reported to be associated with intelligence, the effect sizes were small and almost none of the findings had been replicated. However, since that paper went into press, a number of genetic variants have been shown to be consistently related to intelligence across different studies.

Rietveld et al. (2013)’s meta-analysis found ten SNPs that increased educational attainment, comprising three with nominal genome wide significance and seven with suggestive significance. Recently, a new study has replicated the positive effect of these top three SNPs rs9320913, rs11584700 and rs4851266 on mathematics and reading performance in an independent sample of school children (Ward et al., 2014). Intelligence is a good predictor of performance in educational achievement tests, particularly in subjects such as math and English, where it explained, respectively, 58.6% and 48% of the variance in a longitudinal study based on 70,000+ English children (Deary et al, 2006). Kaufman et al (2012) found high correlations between measures of academic *g* and cognitive *g*. Interestingly, two of the three alleles were among the four with the highest loadings in Piffer's factor analysis of IQ-increasing allele frequencies (Piffer, 2013). For these reasons, I selected only the top three alleles from Rietveld et al.’s meta-analysis whose positive effect on educational attainment was replicated in the study by Ward et al. (2014). Another SNP was selected (rs236330), located within gene *FNBP1L*, whose significant association with general intelligence has been reported in two separate studies (Davies et al, 2011; Benyamin et al, 2013). This gene is strongly expressed in neurons, including hippocampal neurons and developing brains, where it regulates neuronal morphology (Davies et al, 2011).

Principal components analysis (PCA) was used in two separate fashions and with two distinct goals: 1) to detect signals of recent natural selection, or a genotypic intelligence factor, following the method outlined by Piffer (2013), that is selecting trait-increasing alleles and regarding the outcome positive if the extracted component met the following two criteria: a) it represents the first component (explaining the biggest share of the variance); b) all or most of the increasing alleles have high (>0.5) positive loadings on it.

2) To control for genetic distance due to migration (or any statistical artifacts that could arise from analysis of spatial population genetic variation), a random sample of genetic variants (SNPs) is selected, without specifically picking alleles with a particular effect (Tishkoff et al., 2009). The components are then interpreted according to historical information on population movements. The rationale behind this method is that migration acts on all the alleles in the same direction (irrespective of their effect on phenotypes) (Cavalli-Sforza et al., 1996), unlike random drift which unpredictably shifts upwards or downwards allele frequencies at unlinked loci.

Note that although these two uses of PCA appear superficially similar, they produce entirely different results because the former is based on allele pre-selected on the basis of their association with a trait and the factor loadings need to be all (or mostly) positive (if the allele is trait-increasing) for it to be interpretable. The second method, extensively used by Cavalli-Sforza's group, selects random alleles (without regards to their phenotypic effects or possible selection pressure) and does not depend on the sign of factor loadings.

## Methods

IQs were obtained from Lynn (2006) and Lynn & Vanhanen (2012). Some values were updated if they differed markedly from the PISA 2012 results, measuring both fluid intelligence (Creative Problem Solving) and more academic skills (PISA reading, math and science). Finland's IQ was adjusted upwards to 101 (from 97 in Lynn & Vanhanen), to account for recent, more accurate estimates (Armstrong et al., 2014). Spain's IQ was revised downward to 94 (from 97) to accommodate the PISA 2012 results (OECD, 2014). IQ for the Basques was obtained from PISA 2012. IQ for Italy (Tuscany and the north) was obtained from Piffer & Lynn (2014).

Two socio-economic indicators were used: Gross Domestic Product (World Bank, 2014) at purchasing power per capita (GDP (PPP)) and the Human Development Index for 2014 (United Nations Development Programme, 2014).

The frequencies of increaser alleles (those found to increase intelligence or educational attainment in previous GWAS), were obtained from 1000 Genomes (http://www.1000genomes.org/) and ALFRED (http://alfred.med.yale.edu/), returning data (no missing values) for 14 and 50 populations, respectively.

When an SNP was not found on ALFRED, the most closely linked SNPs was searched, and if not found, the second closest and so on, until r^2^≥0.8.Linkage disequilibrium was calculated with SNAP (SNP Annotation and Proxy Search, https://www.broadinstitute.org/mpg/snap/), using the 1000 Genomes pilot 1 dataset, CEU as population panel and a distance limit of 500 kB.

## Results

### 1000 Genomes dataset

A PCA was carried out on the 4 IQ increasing alleles employing the 1K Genomes dataset. A single component (PC) was extracted that explained 77.04% of the variance (the other components had eigenvalues <1 and each explained a small proportion of the variance). Kaiser-Meyer-Olkin (KMO) was acceptable (0.537). Component scores were obtained using the regression method and are reported in table 1 (column 2).

**Table 1*.**

| **Countr**<br>**y** | **IQ PC** | **IQ** | **Predicted**<br>**IQ** | **Z_RES** | **PISA** | **PolHeight\*\*** | **HeightRepl**<br>**\*\*** |
| --- | --- | --- | --- | --- | --- | --- | --- |
| **ASW** | -1.438 | 85 | 76.927 | 1.833 | 422 | 1.34 | 0.65 |
| **LWK** | -1.769 | 74 | 73.372 | 0.142 |  | 1.57 | 0.87 |
| **YRI** | -1.805 | 71 | 72.998 | -0.454 |  | 1.89 | 0.69 |
| **CLM** | -0.136 | 83.5 | 90.865 | -1.673 |  | -0.28 | -0.14 |
| **MXL** | -0.256 | 88 | 89.584 | -0.359 | 420 | -0.4 | 0.08 |
| **PUR** | 0.034 | 83.5 | 92.692 | -2.088 |  | -0.02 | 0 |
| **CHB** | 1.138 | 105 | 104.520 | 0.108 | 577 | -1.35 | -1.69 |
| **CHS** | 0.950 | 105 | 102.507 | 0.566 | 545 | -1.22 | -1.69 |
| **JPT** | 0.931 | 105 | 102.300 | 0.613 | 539 | -1.18 | -1.83 |
| **CEU** | 0.593 | 99 | 98.689 | 0.070 | 520 | -0.11 | 0.93 |
| **FIN** | 0.655 | 100.5 | 99.350 | 0.261 | 543 | -0.34 | 0.59 |
| **GBR** | 0.612 | 100 | 98.886 | 0.253 | 500 | 0.02 | 0.68 |
| **IBS** | 0.003 | 94 | 92.370 | 0.277 | 484 | 0.27 | 0.13 |
| **TSI** | 0.485 | 99.5 | 97.523 | 0.449 | 493 | -0.18 | 0.74 |

\* ASW: African ancestry in SW USA;LWK: Luhya, Kenya; YRI: Yoruba, Nigeria; CLM: Colombian; MXL: Mexican ancestry from LA, California; PUR: Puerto Ricans from Puerto Rica; CHB: Han Chinese in Beijing, China; CHS: Southern Han Chinese; JPT: Japanese in Tokyo, Japan; CEU: Utah Residents with Northern and Western European Ancestry; FIN: Finnish in Finland; GBR: British in England and Scotland; IBS: Iberian population in Spain; TSI: Toscani in Italy.
\*\*PolHeight= PC extracted using 46 alleles (Piffer, 2014). HeightRepl= PC extracted using only the 6 replicated hits (Piffer, 2014).

All four alleles loaded highly (>0.8) and in the right direction (positive sign) on the PC (table 2), indicating that the component represented selection pressure on intelligence or a genotypic intelligence factor (Piffer, 2013).

**Table 2.**
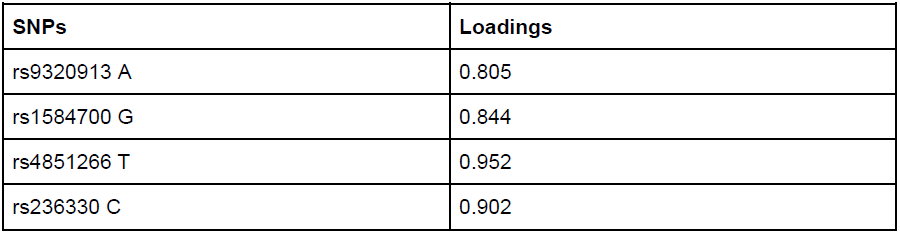
Factor loadings.

| SNPs | Loadings |
| --- | --- |
| rs9320913 A | 0.805 |
| rs1584700 G | 0.844 |
| rs4851266 T | 0.952 |
| rs236330 C | 0.902 |

The correlation between IQ and the PC was very high (r=0.93) and significant (N=14; p=0.000). The correlation between PISA and the PC was strong (r= 0.897; N=10; p= 0.897).

The partial correlation between IQ and the PC after accounting for GDP was highly significant (r=0.937;p=0.000; N=13). A similar result was obtained after partialling out HDI (r=0.942; p=0.000; N=12).

A linear regression (Enter method) was run with IQ as dependent and the PC as independent variable. A significant model emerged (F1,12: 76.49; p=0.000;R²= .865; Adjusted R²= 0.854). Predicted IQ and standardized residuals (Z_RES) were also calculated (table 1, cols. 4–5). The correlation between Z_RES and HDI, GDP were 0.387 and 0.141 but failed to reach significance (p=0.214; p=0.645; N=12 and 13, respectively).

In another regression, Z_RES was used as dependent variable and GDP, HDI were used as predictors to evaluate the effect of socioeconomic factors on phenotypic IQ.

A non-significant model emerged (F2,9: 0.834; p= 0.465; R²= .156; Adjusted R²=−0.031).

Neither HDI nor GDP significantly predicted Z_RES (p=0.791 and 0.794, respectively).

The principal components obtained from 46 alleles (table 1, col.7) and 6 replicated alleles (table 1, col.8) reported in Piffer (2014) were correlated to the IQ PC found in the present study.

The correlations of the IQ PC with “PolHeight” and “HeightRepl” were r=−0.933 (N=14, p= 0.000) and −0.558 (N=14; p=0.038).

To test whether the IQ PC predicted IQ also within races, an univariate ANOVA was carried out with IQ as dependent variable, Group as fixed factor and IQPC as covariate. Four groups were created corresponding to the four continents of 1000 Genomes (Africa=0; America=1; Asia=2; Europe=3).

Both the IQ PC and Group were significant predictors in the model (p=0.032 and 0.029, respectively), indicating that the IQ PC had predictive power also within racial groups (table 3).

**Table 3.**

| Tests of Between-Subjects Effects |  |  |  |  |  |
| --- | --- | --- | --- | --- | --- |
| Dependent Variable: IQ |  |  |  |  |  |
| Source | Type III<br>Sum of<br>Squares | df | Mean<br>Square | F | Sig. |
| Corrected Model | 1634.32<br>1<sup>a</sup> | 4 | 408.580 | 40.99<br>3 | .000 |
| Intercept | 95592.1<br>27 | 1 | 95592.1<br>27 | 9.590,<br>906 | .000 |
| IQPC | 64.071 | 1 | 64.071 | 6.428 | .032 |
| Group | 142.901 | 3 | 47.634 | 4.779 | .029 |
| Error | 89.703 | 9 | 9.967 |  |  |
| Total | 121066.<br>088 | 14 |  |  |  |
| Corrected Total | 1724.02<br>3 | 13 |  |  |  |
| a. R Squared = .948 (Adjusted R Squared = .925) |  |  |  |  |  |

To test whether the IQ PC predicted genotypic height also within races, an univariate ANOVA was carried out with IQ as dependent variable, Group as fixed factor and IQPC as covariate. The results are reported in table 4.

**Table 4.**

| Tests of Between-Subjects Effects |  |  |  |  |  |
| --- | --- | --- | --- | --- | --- |
| Dependent Variable: HeightPol |  |  |  |  |  |
| Source | Type III<br>Sum of<br>Squares | df | Mean<br>Square | F | Sig. |
| Corrected Model | 12.707<sup>a</sup> | 4 | 3.177 | 97.2 | ,000 |
| Intercept | .014 | 1 | .014 | .435 | ,526 |
| IQPC | .156 | 1 | .156 | 4.79 | ,056 |
| Group | 1.393 | 3 | .464 | 14.2 | ,001 |
| Error | .293 | 9 | .033 |  |  |
| Total | 13.000 | 14 |  |  |  |
| Corrected Total | 13.000 | 13 |  |  |  |
| a. R Squared = .977 (Adjusted R Squared = .967) |  |  |  |  |  |

### ALFRED dataset

A PCA was carried out on the 4 IQ increasing alleles (or when these were not available, those in strong linkage disequilibrium: r^2^>0.8) employing the ALFRED dataset.

A single component (PC) was extracted that explained 58.83% of the variance (the other components had eigenvalues <1 and each explained a small proportion of the variance). KMO was good (0.703). All four alleles loaded moderately high on the component (table 6), indicating that it represented selection pressure on intelligence or a genotypic intelligence factor (Piffer, 2013).

**Table 5.**

| **Populations** | **PC1** | **IQ** | **Predicted IQ** | **Z_RES** | **Top4Height** |
| --- | --- | --- | --- | --- | --- |
| **Bantu** | -2.415 | 67 | 67.42 | -0.068 | 2.46 |
| **San** | -2.069 |  | 71.15 |  | 2.42 |
| **Biaka** | -1.634 |  | 75.86 |  | 1.83 |
| **Mbuti** | -1.848 |  | 73.55 |  | 2.02 |
| **Yoruba** | -2.213 | 71 | 69.60 | 0.226 | 2.01 |
| **Mandenka** | -1.896 | 62 | 73.03 | -1.793 | 2.05 |
| **Mozabite** | -0.557 | 84 | 87.49 | -0.568 | 0.45 |
| **Bedouin** | -0.386 |  | 89.34 |  | 0.83 |
| **Druze** | 0.282 | 83 | 96.58 | -2.209 | 0.08 |
| **Palestinian** | 0.153 |  | 95.19 |  | 0.22 |
| **Adygei** | 0.437 |  | 98.26 |  | 0.27 |
| **Basque** | -0.199 | 97 | 91.36 | 0.850 | 0.63 |
| **French** | 0.152 | 100 | 95.17 | 0.784 | 0.55 |
| **CN Italians** | 0.370 | 101 | 97.53 | 0.563 | 0.66 |
| **Orcadian** | 0.881 | 100 | 103.05 | -0.497 | 0.38 |
| **Russian** | -0.345 | 97 | 89.79 | 1.090 | 0.14 |
| **Sardinian** | -0.513 | 94 | 87.97 | 0.978 | 0.34 |
| **Burusho** | 0.302 |  | 96.79 |  | -0.02 |
| **Kalash** | 0.614 |  | 100.16 |  | -0.25 |
| **Pashtun** | -0.239 |  | 90.94 |  | -0.01 |
| **Mongolian** | 1.194 |  | 106.44 |  | 0.14 |
| **Balochi** | 0.244 |  | 96.17 |  | -0.15 |
| **Brahui** | -0.027 |  | 93.23 |  | -0.6 |
| **Hazara** | 0.867 |  | 102.90 |  | -0.17 |
| **Sindhi** | -0.317 | 84 | 90.09 | -0.990 | -0.75 |
| **Dai** | 0.546 |  | 99.43 |  | -0.99 |
| **Daur** | 1.589 |  | 110.71 |  | -0.89 |
| **Han** | 0.947 | 105 | 103.76 | 0.200 | -1.05 |
| **Hezhe** | 1.377 |  | 108.41 |  | -0.88 |
| **Japanese** | 0.725 | 105 | 101.37 | 0.590 | -1.18 |
| **Koreans** | 1.135 | 106 | 105.80 | 0.031 | -1.17 |
| **Lahu** | 0.568 |  | 99.67 |  | -0.94 |
| **Miao** | 0.634 |  | 100.38 |  | -0.99 |
| **Naxi** | 0.210 |  | 95.80 |  | -0.4 |
| **Oroqen** | 0.717 |  | 101.28 |  | -0.68 |
| **She** | 0.416 |  | 98.03 |  | -1.3 |
| **Tu** | 0.726 |  | 101.37 |  | -1.03 |
| **Tujia** | 1.299 |  | 107.57 |  | -0.98 |
| **Uyghur** | 0.805 |  | 102.23 |  | -0.74 |
| **Xibe** | 0.896 |  | 103.21 |  | -0.97 |
| **Yi** | 0.977 |  | 104.09 |  | -0.92 |
| **Cambodians,**<br>**Khmer** | 0.129 | 94 | 94.93 | -0.151 | -0.65 |
| **Papuan New Guinean** | -1.399 | 83 | 78.40 | 0.665 | -0.54 |
| **Melanesian, Nasioi** | -0.225 | 85 | 91.09 | -0.990 | -0.47 |
| **Yakut** | 0.606 |  | 100.08 |  | -0.4 |
| **Pima, Mexico** | -0.284 |  | 90.45 |  | -0.52 |
| **Maya, Yucatan** | -0.945 |  | 83.30 |  | 0.6 |
| **Amerindians** | -1.428 | 86 | 78.08 | 1.28661 | 0.48 |
| **Karitiana** | -0.388 |  | 89.33 |  | 1.04 |
| **Surui** | -0.476 |  | 88.37 |  | 0.01 |

**Table 6.** Factor loadings.

| **SNP** | **Loadings** |
| --- | --- |
| rs9320913 A | .580 |
| rs1584700 G | .820 |
| rs4851266 T | .858 |
| rs236330 C | .780 |

Component scores were obtained using the regression method and are reported in table 5. The correlation between IQ and the PC was very high (r=0.89) and significant (N=19; p=0.000). A linear regression was run (Enter method) with IQ as dependent and the PC as independent variable. A significant model emerged (F1,17: 62.21; p=0.000;R²= .788; Adjusted R²= 0.776). Predicted IQ and standardized residuals (Z_RES) were also calculated (table 5, cols. 4–5).

The correlation between the Height PC and the IQ PC was strongly negative (r= −0.816; p= 0.000; N=50).

To test whether the IQ PC predicted genotypic height also within races, an univariate ANOVA was carried out with IQ as dependent variable, Group as fixed factor and IQPC as covariate.

Seven groups were created (0=African; 1=Middle Eastern; 2=European; 3= Western Asia; 4= East Asian; 5=Native American; 6:South East Asian).

Group was a significant predictor in the model (p= 0.000) but the IQ PC was not (p= 0.768) (table 7) indicating that the inverse correlation between genotypic height and IQ is mediated by racial origin.

**Table 7.**

| **Tests of Between-Subjects Effects** |  |  |  |  |  |
| --- | --- | --- | --- | --- | --- |
| **Dependent Variable: HeightTop4** |  |  |  |  |  |
| **Source** | **Type III Sum of Squares** | **df** | **Mean Square** | **F** | **Sig.** |
| **Corrected Model** | **43.757<sup>a</sup>** | **7** | **6.251** | **53.7** | **.000** |
| **Intercept** | **1.152** | **1** | **1.152** | **9.91** | **.003** |
| **IQPC** | **.010** | **1** | **.010** | **.088** | **.768** |
| **Group2** | **8.360** | **6** | **1.393** | **11.9** | **.000** |
| **Error** | **4.532** | **39** | **.116** |  |  |
| **Total** | **48.332** | **47** |  |  |  |
| **Corrected Total** | **48.290** | **46** |  |  |  |
| **a. R Squared = .906 (Adjusted R Squared = .889)** |  |  |  |  |  |

### Controlling for genetic relatedness

#### 1K Genomes

A random set of 20 SNPs was obtained, each located on a different chromosome (to avoid linkage). These were divided into two sets, belonging to Chr.1-10 and 11–20. Two separate PC's were performed.

The first set produced two components (KMO= 0.676) accounting for 47.78 and 35.89 % of the variance. The second set yielded three components (KMO=0.439), explaining 44.75, 26.88 and 10.9% of the variance respectively.

Table 8 represents the correlation between all the factors. The logic behind this method is that if two factors are not random noise, they should be correlated across sets of different SNPs. Indeed, this is what was found. Factor 1 of set 1 was significantly correlated to factor 1 of set 2; accordingly, factor 2 of set 1 was correlated with factor 2 of set 2. To simplify the analysis (and avoid the issue of multiple comparisons), the two similar pairs were averaged so as to obtain 2 factors (table 9). These were correlated to the variables of interest (IQ, IQ PC). Correlations are reported in table 10.

**Table 8.**

| Correlations |  | FAC1_set2 | FAC2_set2 | FAC3_set2 | fac1set1 | fac2set1 |
| --- | --- | --- | --- | --- | --- | --- |
| FAC1_set2 | Pearson Correlation | 1 | .000 | .000 | .883** | .263 |
|  | Sig. (2-tailed) |  | 1.000 | 1.000 | .000 | .363 |
|  | N | 14 | 14 | 14 | 14 | 14 |
| FAC2_set2 | Pearson Correlation | .000 | 1 | .000 | -.305 | .883** |
|  | Sig. (2-tailed) | 1.000 |  | 1.000 | .289 | .000 |
|  | N | 14 | 14 | 14 | 14 | 14 |
| FAC3_set2 | Pearson Correlation | .000 | .000 | 1 | -.073 | -.290 |
|  | Sig. (2-tailed) | 1.000 | 1.000 |  | .804 | .315 |
|  | N | 14 | 14 | 14 | 14 | 14 |
| fac1set1 | Pearson Correlation | .883** | -.305 | -.073 | 1 | .000 |
|  | Sig. (2-tailed) | .000 | .289 | .804 |  | 1.000 |
|  | N | 14 | 14 | 14 | 14 | 14 |
| fac2set1 | Pearson Correlation | .263 | .883** | -.290 | .000 | 1 |
|  | Sig. (2-tailed) | .363 | .000 | .315 | 1.000 |  |
|  | N | 14 | 14 | 14 | 14 | 14 |
| **. Correlation is significant at the 0.01 level (2-tailed). |  |  |  |  |  |  |

**Table 9.*.** PCs extracted from 20 random SNPs (Averages from the two sets of 10 alleles).

|  | PC1 | PC2 |
| --- | --- | --- |
| ASW | 0.65 | 1.19 |
| LWK | 1.22 | 1.65 |
| YRI | 1.22 | 1.33 |
| CLM | -0.49 | -0.52 |
| MXL | -0.12 | -0.69 |
| PUR | -0.77 | 0.12 |
| CHB | 1.06 | -1.23 |
| CHS | 0.91 | -1.16 |
| JPT | 1.17 | -1.47 |
| CEU | -0.85 | 0.15 |
| FIN | -1.03 | -0.32 |
| GBR | -1.05 | 0.17 |
| IBS | -0.98 | 0.64 |
| TSI | -0.93 | 0.14 |

\* ASW: African ancestry in SW USA; LWK: Luhya, Kenya; YRI: Yoruba, Nigeria; CLM: Colombian; MXL: Mexican ancestry from LA, California; PUR: Puerto Ricans from Puerto Rica; CHB: Han Chinese in Beijing, China; CHS: Southern Han Chinese; JPT: Japanese in Tokyo, Japan; CEU: Utah Residents with Northern and Western European Ancestry; FIN: Finnish in Finland; GBR: British in England and Scotland; IBS: Iberian population in Spain; TSI: Toscani in Italy.

**Table 10.**

| Correlations |  |  |  |  |  |
| --- | --- | --- | --- | --- | --- |
|  |  | IQ | IQPC | AverageFac1 | AverageFac2 |
| IQ | Pearson Correlation | 1 | .930** | -.219 | -.744** |
|  | Sig. (2-tailed) |  | .000 | .451 | .002 |
|  | N | 14 | 14 | 14 | 14 |
| IQPC | Pearson Correlation | .930** | 1 | -.345 | -.839** |
|  | Sig. (2-tailed) | .000 |  | .227 | .000 |
|  | N | 14 | 14 | 14 | 14 |
| AverageFac1 | Pearson Correlation | -.219 | -.345 | 1 | -.011 |
|  | Sig. (2-tailed) | .451 | .227 |  | .970 |
|  | N | 14 | 14 | 14 | 14 |
| AverageFac2 | Pearson Correlation | -.744** | -.839** | -.011 | 1 |
|  | Sig. (2-tailed) | .002 | .000 | .970 |  |
|  | N | 14 | 14 | 14 | 14 |
| **. Correlation is significant at the 0.01 level (2-tailed). |  |  |  |  |  |

Factor 1 was not significantly correlated either to IQ or IQ PC and was not clearly interpretable, but perhaps it indicated north-south migrations. Factor 2 had significantly negative correlations with both. This factor likely represents genetic relatedness to Africans (due to migration), as it has higher values among Africans and lower among Europeans, lower still among Americans and the lowest among East Asians, similar to genetic distances reported using other genetic markers (Cavalli-Sforza et al., 1996). It's a common finding from PCA of genetic markers that the difference between Africans and non-Africans is greater than between other populations, as reflected by the first PC usually distinguishing between Africans and non-Africans in worldwide samples (Tishkoff et al., 2009).

A partial correlation was carried out to assess the relationship between IQ and the estimated genotypic IQ factor (IQ PC), partialling out the “African” factor (Factor 2).

The partial correlation was still highly significant (r= 0.841; p=0.000; N=14). As a cross-check, another partial correlation was computed between the “African” factor (Factor 2) and IQ, after partialling out IQ PC. The rationale is that if the IQ PC really represents an intelligence factor due to selective pressure and is not a statistical artefact or a by-product of migrations, then partialling out non-selective factors (such as African admixture) will not substantially alter its effect on IQ. Conversely, if the African factor does not represent a genuine intelligence factor due to selection, its correlation with IQ will be spurious (i.e. it will disappear after partialling out the IQ PC). This is indeed what was found: the partial correlation between IQ and Factor 2 (“African”) after controlling for IQ PC became negligible and even reversed its sign (r= 0.181; p= 0.554).

To assess the effects of all genetic factors, a multiple linear regression was ran with IQ as dependent variable and IQ PC, PCs 1,2 as predictors. Results are reported in table 11. IQ PC emerged as the only significant predictor of IQ, with a SD change corresponding to an increase of over 1 SD in population IQ. PC 2 lost all its negative association with IQ and even had a small positive effect, albeit failing to reach significance. This suggests that its negative correlation with IQ is spurious (i.e. mediated by the intelligence factor).

**Table 11.**

| Coefficients |  |  |  |  |  |  |
| --- | --- | --- | --- | --- | --- | --- |
| Model |  | Unstandardized Coefficients |  | Standard Coefficient | t | Sig. |

|  |  | B | Std. Error | Beta |  |  |
| --- | --- | --- | --- | --- | --- | --- |
| 1 | (Constant) | 92.328 | 1.096 |  | 84.250 | .000 |
|  | IQPC | 15.184 | 2.752 | 1.319 | 5.517 | .000 |
|  | AverageFac1 | 2.838 | 1.543 | .239 | 1.839 | .096 |
|  | AverageFac2 | 4.328 | 2.663 | .365 | 1.626 | .135 |
| a. Dependent Variable: IQ |  |  |  |  |  |  |

Another multiple regression was run with IQ PC as dependent and the genotypic height factor (PolHeight) plus the two migratory factors. Again genotypic height was the only significant predictor (Beta= −1.44; p= 0.02).

#### ALFRED

A set of 10 SNPs (different from those used in the 1K analysis) was randomly chosen (Chr.1-10). Principal component analysis extracted three components with eigenvalues greater than 1, accounting for 32.86, 16.99, 14.6 % of the variance. KMO was acceptable (0.608). PC scores are reported in table 12. The first component clearly represents genetic distance from Africans, as African populations show the highest values, and the component scores for the other populations follow the well-established pattern of progressive distance from Africans in the following order (found in studies employing much larger numbers of genetic markers): Middle Eastern> European>SW Asian>East Asians>Native American>Oceanian (Eurogenes, 2014). The other components had no clear interpretation and since they accounted for a small portion of the variance, it is likely that they constitute statistical noise.

**Table 12.** “Migration” PCs from 10 SNPs (ALFRED).

|  | PC1 | PC2 | PC3 |
| --- | --- | --- | --- |
| **Bantu** | 1.946 | -0.240 | 1.184 |
| **San** | 1.840 | 0.236 | -0.513 |
| **Biaka** | 2.869 | -0.179 | 2.401 |
| **Mbuti** | 1.929 | -0.439 | 0.000 |
| **Yoruba** | 1.997 | 0.035 | 2.019 |
| **Mandenka** | 1.683 | 0.041 | 1.435 |
| **Mozabite** | 0.613 | 0.798 | 0.333 |
| **Bedouin** | 0.812 | 1.452 | -0.149 |
| **Druze** | 0.409 | 1.275 | -0.783 |
| **Palestinian** | 0,673 | 1.474 | -0.529 |
| **Adygei** | -0.174 | 2.229 | -0.615 |
| **Basque** | -0.518 | 0.855 | -0.430 |
| **French** | 0.173 | 1.447 | -0.592 |
| **CNIItalians** | -0.010 | 1.181 | -0.924 |
| **Orcadian** | -0.635 | 1.894 | -0.063 |
| **Russian** | -0.137 | 0.942 | -0.558 |
| **Sardinian** | 0.011 | 1.394 | -1.080 |
| **Burusho** | 0.403 | -0.243 | -0.888 |
| **Kalash** | 0.257 | -0.149 | 0.068 |
| **Pashtun** | 0.222 | -0.573 | -0.508 |
| **Mongolian** | -0.688 | -0.387 | -0.215 |
| **Balochi** | 0.223 | 0.643 | -0.502 |
| **Brahui** | 0.380 | -0.071 | -0.605 |
| **Hazara** | -0.434 | 0.290 | -0.163 |
| **Sindhi** | -0.265 | 0.027 | 0.051 |
| **Dai** | 0.395 | -2.021 | -0.846 |
| **Daur** | -0.976 | -0.124 | -0.241 |
| **Han** | -0.191 | -0.900 | -0.733 |
| **Hezhe** | -0.415 | 0.042 | 0.652 |
| **Japanese** | -0.376 | -1.672 | -0.265 |
| **Koreans** | -0.040 | -0.995 | -0.212 |
| **Lahu** | -0.224 | -1.065 | 0.040 |
| **Miao** | -0.457 | -0.385 | -0.673 |
| **Naxi** | 0.311 | -0.716 | -1.521 |
| **Oroqen** | -0.564 | 0.091 | -0.118 |
| **She** | -0.248 | -1.092 | -0.594 |
| **Tu** | 0.358 | -0.584 | -0.543 |
| **Tujia** | 0.111 | -1.667 | -0.841 |
| **Uyghur** | 0.267 | 0.352 | 0.052 |
| **Xibe** | -1.228 | -0.285 | -0.661 |
| **Yi** | 0.348 | -0.9634 | -0.220 |
| **Cambodians, Khmer** | 0.021 | -1.681 | -0.651 |
| **Papuan New Guinean** | -1.555 | -0.496 | -0.384 |
| **Melanesian, Nasioi** | -1.044 | -0.352 | -0.379 |
| **Yakut** | -0.795 | 0.361 | 0.069 |
| **Pima, Mexico** | -1.293 | -0.550 | 1.931 |
| **Maya, Yucatan** | -1.004 | 0.043 | 0.477 |
| **Amerindians** | -1.224 | -0.251 | 2.219 |
| **Karitiana** | -1.807 | -1.135 | 2.995 |
| **Surui** | -1.948 | 2.117 | 2.081 |

The correlation between the first component and the genotypic IQ factor was negative (r=−0.468, N=50, p=0.000).

A multiple linear regression was run with IQ as dependent variable and genotypic IQ factor plus the three “migration components”. A significant model emerged (F4,14= 15.77, p=0.000; (R²=.818; Adjusted R²= 0.767). Only the genotypic IQ factor was a significant predictor of IQ. Beta coefficients and p values are reported in table 13.

**Table 13.**

| **Coefficients<sup>a</sup>** |  |  |  |  |  |  |
| --- | --- | --- | --- | --- | --- | --- |
| **Model** |  | **Unst.Coeff.** |  | **St.Coeff.** | **t** | **Sig.** |
|  |  | **B** | **S.E.** | **Beta** |  |  |
| **1** | **(Constant)** | 93.180 | 1.622 |  | 57.44 | .000 |
|  | **IQPC** | 9.491 | 2.125 | .780 | 4.465 | .001 |
|  | **PC1** | -2.594 | 1.751 | -.195 | -1.481 | .161 |
|  | **PC2** | -.048 | 1.443 | -.004 | .,033 | .974 |
|  | **PC3** | -.286 | 2.295 | -.022 | -.124 | .903 |
| **a. Dependent Variable: IQ** |  |  |  |  |  |  |

Another multiple regression was run with IQ PC as dependent and the genotypic height factor plus the three migratory principal components. IQ PC had a significantly negative effect (Beta= −0.838; p= 0.000) but the other variables did not achieve significance at the p= 0.05 level.

## Discussion

The genotypic IQ factor extracted with principal component analysis was a strong predictor of average population IQ, almost approaching unity in the 1K Genomes dataset and showing strong effects in the ALFRED dataset, comprising only partly overlapping populations.

An additional analysis revealed that this effect was not entirely mediated by race, because the genotypic factor significantly predicted IQ in both datasets (1K Genomes and ALFRED) consisting of four and seven racial groups, respectively (see table 3 and 7). If replicated, this would be a remarkable finding, which implies that evolutionary forces persisted even after continental races split into populations that roughly correspond to contemporary national or ethnic groups. This also yields more credibility to the results, because it indicates that the effect is not due to a broader racial element but is repeated across isolated geographical regions and genetic clusters.

For instance in Europe, northern populations (Finland, Great Britain) have higher genotypic scores than southern populations (Iberians), which could be due to lower admixture with Africans or higher selective pressure for intelligence. This corresponds to an advantage in terms of predicted IQ of almost 6 points (table 1, col. 4), which is reflected in the phenotypic IQ (col. 2).

The admixture scenario is unlikely, as Iberians have only low (about 2%) frequency of sub-Saharan genetic markers (Eurogenes, 2014).

However, the higher genotypic score of African Americans compared to sub-Saharan Africans (77 vs 73) is obviously due to admixture with people of European ancestry. The difference in phenotypic IQ between them is much higher (10–15 points), which is probably due to the effect of malnutrition that tends to depress intellectual development. It is thus likely that the real sub-Saharan African genotypic IQ is higher than estimated here or what is measured and instead is around 81 (85-4).

An attempt was made to determine whether the discrepancy between phenotypic and genotypic IQ was due to socioeconomic factors. These (GDP per capita, Human Development Index) failed to predict the residuals when entered in a regression. It should also be mentioned that although they failed to reach significance, the correlations between residuals and GDP/HDI were in the right direction, that is to say, larger residuals (phenotypic-genotypic IQ) were associated with higher GDP and better human development. The lack of statistical significance is most likely due to the small sample size (N<15).

When allele frequency data will become available for more populations, it will be possible to test this hypothesis on a bigger sample. The populations from the ALFRED database comprise in part tribes (Papuans, Melanesian, Amerindians) and assigning GDP or HDI to them was not possible. Also carrying out a between-groups analysis was problematic because IQ scores for some racial groups were available only for a single population.

However, the ALFRED results also yield interesting insights.

The populations with the highest genotypic IQ are the Daur and Hezhe (or Nanai) (table 5, col. 4).

The Daur speak a Mongolic language and are descendants of the Khitan (Jinhu, 2001), who were originally from Mongolia and Manchuria (the northeastern region of modern-day China). The Hezhe are a Tungusic people who have traditionally lived in northeastern China and Siberia.

This finding (if replicated) would yield support to the “cold winters theory”, which posits that the higher IQ of East Asians originated in a region roughly corresponding to modern day Mongolia and Manchuria, characterized by extremely low winters temperatures and then spread to the south of China and Japan through migrational waves (Lynn, 1987). There is recent genetic evidence that fits well with a model of north-south migration in China: “The inferred north-south pattern in the genetic structure analyses suggests a primary north-south migratory pattern in China. This ties in very well with historical records indicating that the *Huaxia* tribes in northern China, the ancient ancestors of the Han Chinese, embarked on a long period of continuous southward expansion as a result of war and famine over the past two millennia" and “this one-dimensional structure of the Han Chinese population is clearly characterized by a continuous genetic gradient along a north-south geographical axis, rather than a distinct clustering of northern and southern samples" (Chen et al., 2009).

Some populations that ALFRED groups with East Asians (because they live there) are in fact descended from South East Asians, for example the Lahu, the Naxi and the Dai, and have a lower genotypic IQ (95–99) than other groups living in China (100–110), resembling the genotypic IQ of Cambodians (95) (Table 5).

Two observations deserve attention, regarding a marked discrepancy between genetic distance (as measured by neutral genomic markers) and distance in factor scores for IQ and height.

(1) The lower IQ of Sub-Saharan Africans and the Aboriginals from Papua New Guinea cannot be explained in terms of cultural contact or genetic relatedness, because these racial groups are geographically very far apart and genetically very distant from each other (Cavalli-Sforza et al., 1996).
(2) Native Americans have very different genotypic IQ and height from East Asians, being taller and with lower phenotypic and genotypic IQ. This contrasts markedly with their genetic similarity and relatively recent common origin (Wells, 2002) and argues strongly against the possibility that random drift can explain this fact, because the increase in height and decrease in IQ follows the worldwide pattern found in the present study. Instead it lends support to a model of an evolutionary trade-off between increased height or higher intelligence.

These two variables were strongly negatively related in both datasets (1K Genomes and ALFRED).

The negative relationship between genotypic height and IQ seems to be mediated by race, as the IQ PC failed to predict height after partialling out the effects of race (group). This suggests that the evolutionary trade-off between height and IQ predates sub-continent level population splits, although this does not rule out that it continued after (as suggested by the marginally significant independent effect of genotypic IQ on genotypic height in the 1K Genomes dataset). It can only be speculated that this trade-off was due to sexual selection (for brawny vs brainy males) or selective pressure due to climate, favoring shorter limbs (via Allen's rule) and bigger brains in colder environments (Piffer, 2014). This between-groups pattern contrasts with the within group pattern, where IQ is slightly higher on average in taller people than in shorter people (Pearce et al., 2005; Humphreys, Davey & Park, 1985). This correlation is in part due to common genetic factors (Marioni et al., 2014) and assortative mating (Beauchamp et al., 2011). It is possible that mutational load has decreased both traits, hence creating the genetic correlation, which would be strengthened by assortative mating and environmental variables (e.g. nutrition).

A set of randomly picked SNPs located throughout the genome was used to estimate genetic relatedness between populations due to migrations or possible statistical artifacts. This methodology was first used by Cavalli-Sforza and colleagues over 30 years ago to study the evolutionary history of human populations and reconstruct patterns of migration, and it continues to be used today (Ma & Amos, 2010).

If the component extracted from the IQ increasing alleles really reflects selection pressures on intelligence, it will predict IQ also after partialling out the confounding element due to population migrations. Although the biggest migration principal component (indicating genetic distance from Africans) was negatively correlated to IQ, the genotypic IQ factor emerged as a significant independent predictor of IQ in both datasets (ALFRED and 1000 Genomes), independently of the other (migration+noise) components.

This analysis revealed that the effect on IQ of the genotypic IQ factor is stronger than migration and is not substantially mediated by it.

A similar analysis showed that the genotypic height factor had a significantly negative and strong effect on the genotypic IQ factor even accounting for the migratory PCs, in both datasets (ALFRED and 1K Genomes).

Overall the results provide support for the validity of the method proposed by Piffer (2013) for detecting signals of recent polygenic selection and indicate that the principal components that were extracted are clearly distinguishable from those produced by migrations.

A clear limitation of this study is its reliance on a very small number of genetic variants for intelligence (4) although the positive results obtained with a much bigger sample of alleles (N=46) affecting height (Piffer, 2014) are encouraging. Future studies using more genetic variants will have to attempt a replication of the findings presented here.

